# Effects of parental age and polymer composition on short tandem repeat *de novo* mutation rates

**DOI:** 10.1101/2023.12.22.573131

**Authors:** Michael E. Goldberg, Michelle D. Noyes, Evan E. Eichler, Aaron R. Quinlan, Kelley Harris

**Author notes:** These authors contributed equally to this work (Aaron R. Quinlan, Kelley Harris).

## Abstract

Short tandem repeats (STRs) are hotspots of genomic variability in the human germline because of their high mutation rates, which have long been attributed largely to polymerase slippage during DNA replication. This model suggests that STR mutation rates should scale linearly with a father’s age, as progenitor cells continually divide after puberty. In contrast, it suggests that STR mutation rates should not scale with a mother’s age at her child’s conception, since oocytes spend a mother’s reproductive years arrested in meiosis II and undergo a fixed number of cell divisions that are independent of the age at ovulation. Yet, mirroring recent findings, we find that STR mutation rates covary with paternal and maternal age, implying that some STR mutations are caused by DNA damage in quiescent cells rather than the classical mechanism of polymerase slippage in replicating progenitor cells. These results also echo the recent finding that DNA damage in quiescent oocytes is a significant source of *de novo* SNVs and corroborate evidence of STR expansion in postmitotic cells. However, we find that the maternal age effect is not confined to previously discovered hotspots of oocyte mutagenesis, nor are post-zygotic mutations likely to contribute significantly. STR nucleotide composition demonstrates divergent effects on DNM rates between sexes. Unlike the paternal lineage, maternally derived DNMs at A/T STRs display a significantly greater association with maternal age than DNMs at GC-containing STRs. These observations may suggest the mechanism and developmental timing of certain STR mutations and are especially surprising considering the prior belief in replication slippage as the dominant mechanism of STR mutagenesis.

**Author Summary:** We have long known that tandem repeats are hypermutable and attributed that hypermutability to slippage during DNA replication. Contradicting this long-held theory, we show that tandem repeats accumulate mutations in maternal germ cells during periods when these cells do not replicate. This bolsters a new consensus that DNA replication is not the only driver of mutagenesis, even at loci where replicative slippage is possible. Patterns shared by certain loci enriched for mutations from older mothers may hint at mechanisms.

## Introduction

Short tandem repeats (STRs) are genomic elements composed of repeating nucleotide motifs typically of length 1-6bp (Weber and Wong 1993). Sometimes also known as micro- and minisatellites, they mutate at rates orders of magnitude greater than non-repetitive loci, a feature which results in high levels of diversity that enabled studies of genetic diversity in the pre-genomic era (Weber and Wong 1993; Spencer *et al*. 2000). Despite this classical importance, STRs have fallen behind relative to single nucleotide variants (SNVs) in terms of our understanding of how germline mutation rates vary in humans, what environmental or genetic conditions may contribute to this variance, and how their variation broadly shapes complex traits and genomic evolution.

STRs are numerous – they encompass nearly 3% of the human genome, a similar proportion to that of coding regions (International Human Genome Sequencing Consortium *et al*. 2001). Until recently, our understanding of how specific STR loci affect traits remained limited to a small number of large effect size loci, though the molecular underpinnings of their associations still remain largely unclear (McGinty and Sunyaev 2023). Germline and somatic expansion at these loci have long been implicated in Mendelian developmental and neurodegenerative disorders (Sherman *et al*. 1985; Gatchel and Zoghbi 2005; Mitra *et al*. 2021; Nurk *et al*. 2022). More broadly, variation genome-wide has recently demonstrated association with complex traits and gene expression (Gymrek *et al*. 2016; Fotsing *et al*. 2019; Margoliash *et al*. 2022). Accordingly, some STRs are inferred to be under purifying selection, regardless of their instability (Gymrek *et al*. 2017; Mitra *et al*. 2021). Children diagnosed with simplex (likely non-inherited) Autism Spectrum Disorder (ASD) have been found to harbor *de novo* mutations (DNMs) at STRs inferred to be more deleterious than those identified in their unaffected siblings (Mitra *et al*. 2021). These recent findings suggest that STRs’ complex genotype-phenotype interactions and evolutionary history may have similar complexity to those attributed to SNVs and structural variation.

Both SNV and STR mutation rates depend strongly on paternal age (Kong *et al*. 2012; Forster *et al*. 2015). This dependency was classically explained by the fact that male germ cells replicate throughout the reproductive lifespan, assuming that most mutations originate as errors in replication (Goodman 1985; Drost and Lee 1995). In the case of SNVs, recent work has cast doubt on this model and has begun to support DNA damage as a substantial source of SNVs (Goodman 1985; Drost and Lee 1995; Gao *et al*. 2019; Wu *et al*. 2020). The proportion of SNVs inherited from the paternal lineage, known as *ɑ*, remains strikingly stable with parental age, which should not be the case if most mutations are replicative in origin and the ratio of paternal to maternal cell divisions increases with age (Gao *et al*. 2019). Although the female germline does not continue to replicate after birth, the ratio of male to female germline mutations appear to remain largely constant regardless of developmental stage. Nevertheless, analyzing how the rates of different types of mutations covary with parental age and sex can provide clues to mutational mechanisms that may be more associated with replication errors or with DNA damage.

Although we have less information on how life history impacts STR mutagenesis, the high mutation rate at these loci has long been attributed primarily to polymerase slippage during S phase replication (Klintschar *et al*. 2004; Forster *et al*. 2015). Replication slippage typically causes STR expansions or deletions by one or more repeat units (Brinkmann *et al*. 1998; Sun *et al*. 2012; Amos *et al*. 2015; Mitra *et al*. 2021). Historically, studies have found no genome-wide association of STR mutation rates with maternal age (maternal age effect) but have found a strong association with paternal age (paternal age effect) (Sun *et al*. 2012; Forster *et al*. 2015). At face value, these age effects seemed to bolster the theory that STRs mutate only during DNA replication in S phase. Given the lack of maternal germline replication after the future mother’s birth, we would expect replication-driven mutations to exhibit a paternal age effect but no maternal age effect (Drost and Lee 1995). However, previous studies of STR parental age associations draw upon much less data than comparable studies of single nucleotide mutation rates. Certain STR loci have been shown to accumulate mutations in post-mitotic cells, indicating that damage could play some role in STR mutagenesis independent of cell divisions (Gonitel *et al*. 2008). For example, somatic expansion in postmitotic striatal medium spiny neurons of the CAG/CAA repeat that encodes a polyglutamine tract in the *huntingtin* gene causes Huntington’s Disease, a highly heritable and penetrant neurodegenerative disease with adolescent/early adulthood onset (Wright *et al*. 2019; Lee *et al*. 2019). Furthermore, *in vitro* studies have shown evidence that the rate of expansions and deletions at STRs may not be fully explainable by strand slippage alone (Ananda *et al*. 2011).

Although directly proving a link between a mutational pattern and its causal mutational pathway can be challenging, it is possible to infer potential molecular mechanism by examining the covariance of mutation rate with polymer complexity and nucleotide content. Local mutation rate, chromatin state, and many other genomic features covary with nucleotide content (Hwang and Green 2004; Alexandrov *et al*. 2013, 2020; Goldberg and Harris 2022). This broad phenomenon can be attributed to the fact that nucleotide content is highly predictive of local DNA conformation, or 3-dimensional molecular shape, which in turn associates with the types of molecules likely to interact with a locus and the frequency of those interactions (Rohs *et al*. 2009; Lazarovici *et al*. 2013; Abe *et al*. 2015; Liu and Samee 2023). Similarly, certain types of STRs are known to form unique structures different from the canonical double helix of B DNA; for example, some guanine rich repeats form fragile cruciforms (McGinty and Sunyaev 2023). Different tumor types display variance in mutational spectra and signatures in mutations at repetitive DNA elements, indicating tissue- or tumor-specific mutation rate modifiers (Alexandrov *et al*. 2020). When applied to polymorphic and *de novo* germline SNVs, these types of analyses have revealed significant diversity in the spectra of mutations accumulating in different populations, families, or species (Hwang and Green 2004; Harris 2015; Harris and Pritchard 2017; Carlson *et al*. 2020; Bergeron *et al*. 2023). Similar approaches have helped identify variance in mutational pathways active at STRs: recently, a small set of families were discovered to harbor a mutator allele that affects indels at TNT repeats (Reijns *et al*. 2022; Maksimov *et al*. 2023).

In this study, we sought to further describe how STR mutation rates vary as a function of parental age and sex to better understand the developmental timeline and possible mutational mechanisms. To do so, we analyzed a dataset of *de novo* mutations (DNMs) at STR loci found in 1593 families comprising two parents and two children (quads), one of whom has been diagnosed with a simplex case of ASD without prior family history (Fischbach and Lord 2010). These families were recruited, in part, to study how rare and *de novo* mutations contribute to ASD as a subset of the Simons Simplex Collection (SSC) (Fischbach and Lord 2010). Our work aims to clarify how parental age and sex impact both autism risk and deleterious DNM load, as we further characterize how these variables effect the distribution of fitness effects of STR DNMs. Very recently, a study examining nearly 77k STR DNMs in over 6000 trios identified a significant association of DNM rate with maternal age, dispelling a simplistic model of STR mutagenesis in the germline (Kristmundsdottir *et al*. 2023). We confirm the presence of this maternal age effect in an independent dataset and demonstrate its robustness to noise typical of STRs genotyped with short read sequencing. Furthermore, we examine how parental age effects are modified by certain genomic features, identifying specific DNA motifs enriched for the maternal age signal.

## Results

We examined STR *de novo* mutations in a cohort of 1593 quad families recruited as a part of the Simons Simplex collection based on the diagnosis of a simplex case of ASD in a single child (Fischbach and Lord 2010). These probands have been shown to harbor an enrichment of both SNV and STR DNMs that may affect neurodevelopment and lead to ASD (Turner *et al*. 2017; Mitra *et al*. 2021). We used mutation calls from Mitra et al., 2021. As described previously, the families were sequenced to 35X coverage, and diploid genotypes at STR loci were inferred using GangSTR (Fischbach and Lord 2010; Mitra *et al*. 2021). This database of STR DNMs is the largest compiled to date. DNMs were called using MonSTR, a likelihood-based caller with an accuracy of 90%; this accuracy was estimated based on validation with capillary electrophoresis (Mitra *et al*. 2021). Stringent filters were placed on the genotype calls as described in Mitra et al., 2021. We developed further stringent filters on the DNM calls to account for allelic dropout in parents at high diversity sites that result in false positive DNM calls in the children; these filters were further validated with Pacific Biosciences high fidelity (HiFi) and Oxford Nanopore Technologies long read sequencing generated for two quads (Methods) (Figures S1, S2; Table S1). Briefly, these filters involve removing any DNM at a locus at which one or more reads observed in the parents matches the putative *de novo* allele observed in the child and resulted in filtering out 43% of non-homopolymer DNMs (30623 of out 71581). Similarly to Mitra et al., we excluded homopolymers from all analyses unless otherwise specified, as homopolymers are challenging to genotype with short read next generation sequencing methods (Stoler and Nekrutenko 2021). As fully described in Mitra et al., mutations were phased to their parental lineages based solely on the observed alleles in parents and child without explicitly recreating the surrounding haplotype. If the inherited allele in a child could be assigned unambiguously to one parent’s lineage, the *de novo* allele was phased to the other parent’s lineage (Mitra *et al*. 2021). We were able to phase 56% of the non-homopolymer DNMs that passed our filters. Following these filtering steps, the mean number of STR DNMs per child was 12.91; the mean number of paternally and maternally phased DNMs was 5.80 and 1.43, respectively (95% confidence intervals [CI] 12.76 – 13.06, 5.70 – 5.89, and 1.39 - 1.49, respectively). Affected probands and their unaffected siblings respectively harbored 13.11 and 12.72 STR DNMs (95% CI 12.96 - 13.25, 12.57 - 12.87, respectively).

**Figure S1:**
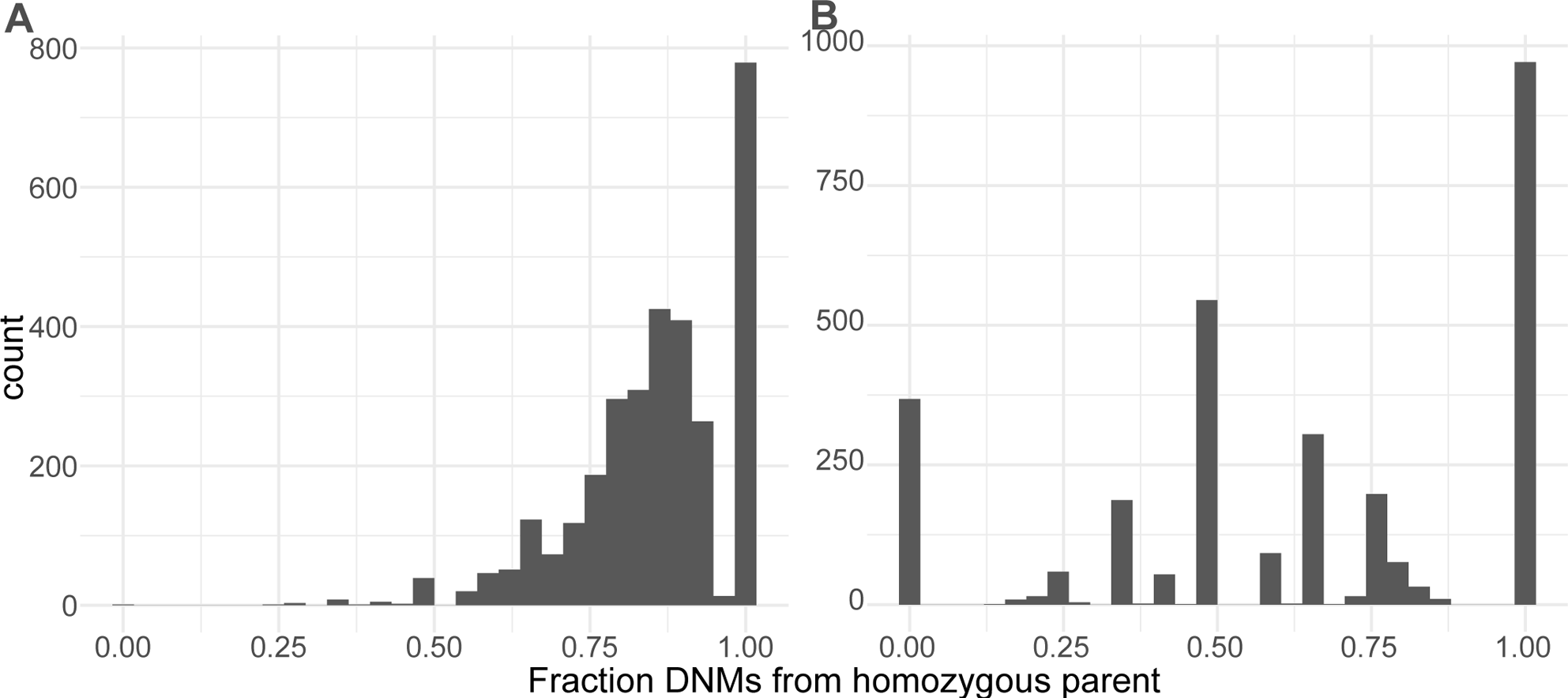
Inheritance bias potentially supports allelic dropout. (A) We observed that, prior to filtering, DNMs at loci where one parent was heterozygous and the other was homozygous nearly always phased to the homozygous parent. Here we have plotted the fraction of mutations per child at those types of sites that phase to the homozygous parent. (B) Once we filter out DNMs where ≥ 1 read in the parent contains the putative *de novo* allele, we observe a reduction in the skew towards inheritance from the homozygous parent. The behavior following this filtering supports allelic dropout as a possible erroneous source of DNMs.

**Figure S2:**
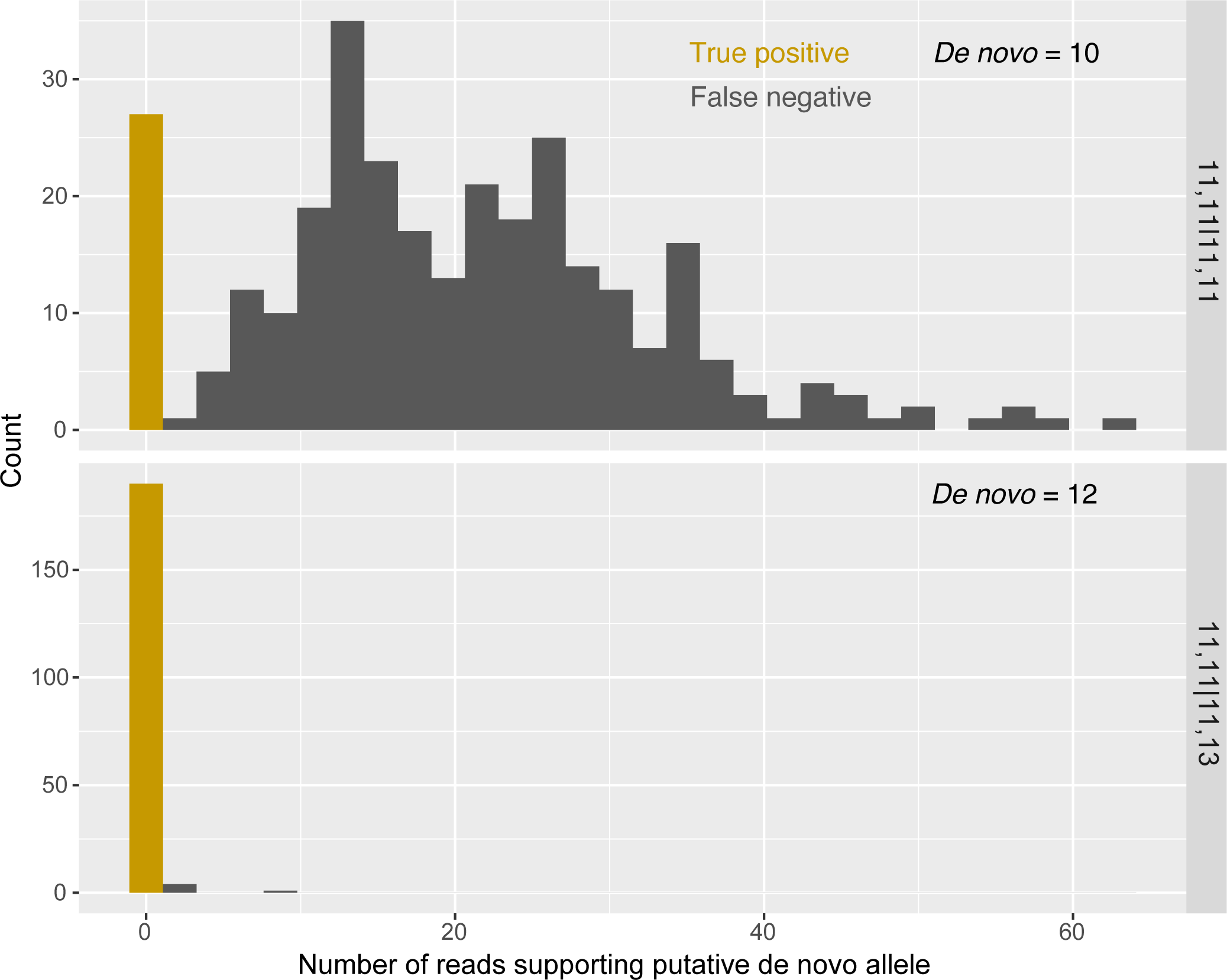
Variance in filtration sensitivity between genotypes at the same locus (chr22: 27079676). We filter out any STR DNM when >0 reads across either parent support the putative *de novo* allele. This filtering strategy is strict; we therefore explored the effect on filter sensitivity of a allowing a higher threshold of parental reads supporting the de novo allele. Putative *de novo* alleles were identified in children of two different families at the same locus; parental and child genotypes in the first family were [11,11], [11,11], and [11,10], respectively; the second family were [11,11], [11,13], and [11,12], respectively. To estimate sensitivity of our filtration strategy, we identified a set of “positive” couples with identical genotypes to the targeted family but no putative *de novo* allele in their children; we believe that allelic dropout is unlikely to have occurred when genotyping these positive couples. To assess how frequently we would erroneously flag these couples’ genotypes for allelic dropout and explore whether to allow a higher threshold of reads mapping to a putative *de novo* allele. Plotted in the histograms are the distribution of the number of reads, summed between both parents, that support the putative *de novo* allele in the positive couples. Filter sensitivity is higher for couples with [11,11] and [11,13] than for couples with [11, 11] and [11,11].

We separately regressed the number of maternally and paternally phased mutations per child against their mother’s and father’s ages at their birth, respectively, using Poisson regressions with identity link functions (this regression model was used throughout the study unless otherwise specified). Our analysis confirms the strong paternal age effect on the rate of paternally phased STR mutations that has been reported in this and other STR DNM datasets (Figure 1a) (0.14 paternal mutations/year, std. err. = 8.01 * 10^−3^, *P* = 1.11*10^−66^). In line with other recent findings, we also observe a significant effect of maternal age on maternally inherited STR DNMs (Figure 1b) (0.025 maternal mutations/year, std. err. = 4.42 * 10^−3^, *P* = 8.11*10^−9^) (Kristmundsdottir *et al*. 2023). The maternal age model does not demonstrate significant overdispersion, which can lead to artificially deflated *P* values (*P* = 0.27, dispersion = 1.02, overdispersion test). Maternal age remains a significant predictor of the maternal STR mutation rate even when we include paternal age as a covariate, indicating that phasing errors are not likely to explain the signal (*P* = 4.20*10^−6^). An exponential model of the maternal age effect fits slightly better than the linear model (delta AIC = −0.2), whereas the linear model is the superior fit for the paternal age effect (delta AIC = −5). This finding mirrors the observation of an exponential maternal age effect on SNV rate presented in Gao et al., 2019 (Gao *et al*. 2019). Interestingly, neither paternal nor maternal age was significantly associated with the DNM rate at homopolymers (*P* = 0.062, 0.816 for paternal and maternal age effects, respectively) (Figure S3). Although this observation could support an association between parental age effects and variance in a mutational pathway that does not affect homopolymers, it may also reflect the difficulty of accurately genotyping homopolymers and that any parental age effects are overwhelmed by the noise of sequencing errors (Stoler and Nekrutenko 2021). We replicated these analysis accounting for possible additional bias in genotyping and still observed a significant association between maternal age and DNM rate (Methods).

**Figure 1:**
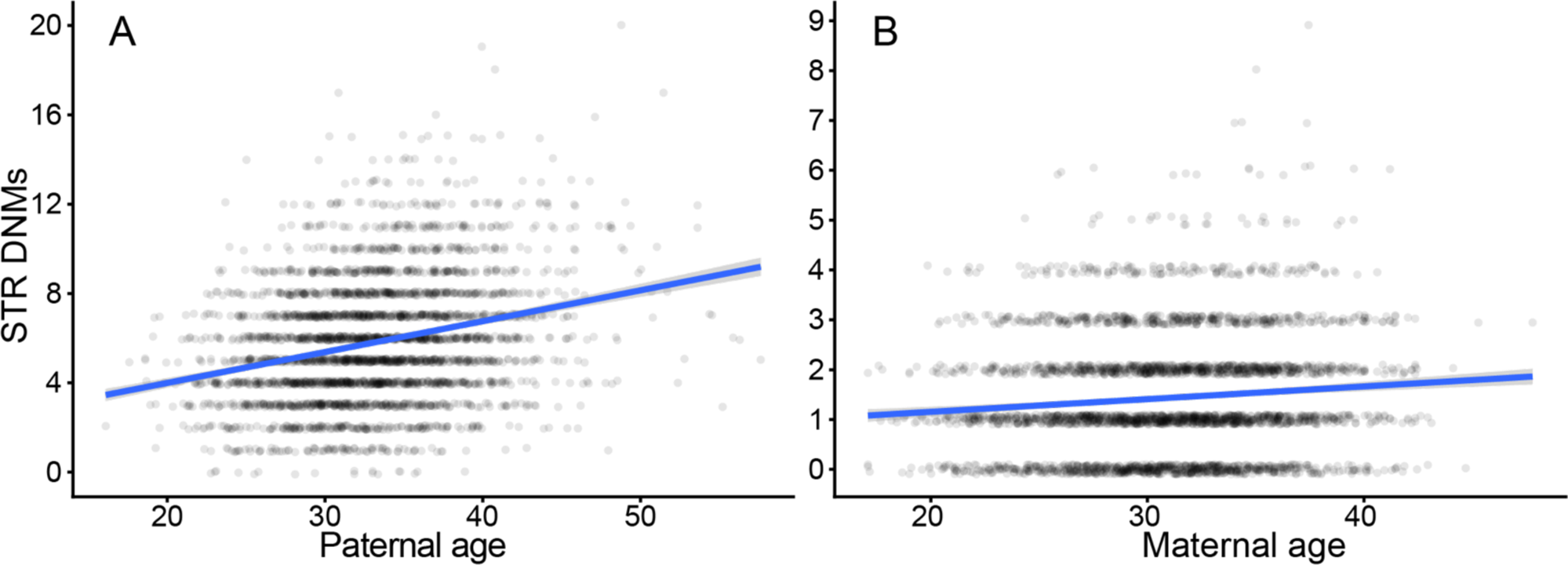
Significant maternal and paternal age effects on STR DNM rate. A, B: Number of paternal and maternal nonhomopolymer STR DNMs per offspring in the SSC are plotted and regressed against paternal and maternal age at birth, respectively. Y-axis jitter is added to points for readability. The regression line is a Poisson GLM fit with an identity link function in ggplot2.

**Figure S3:**
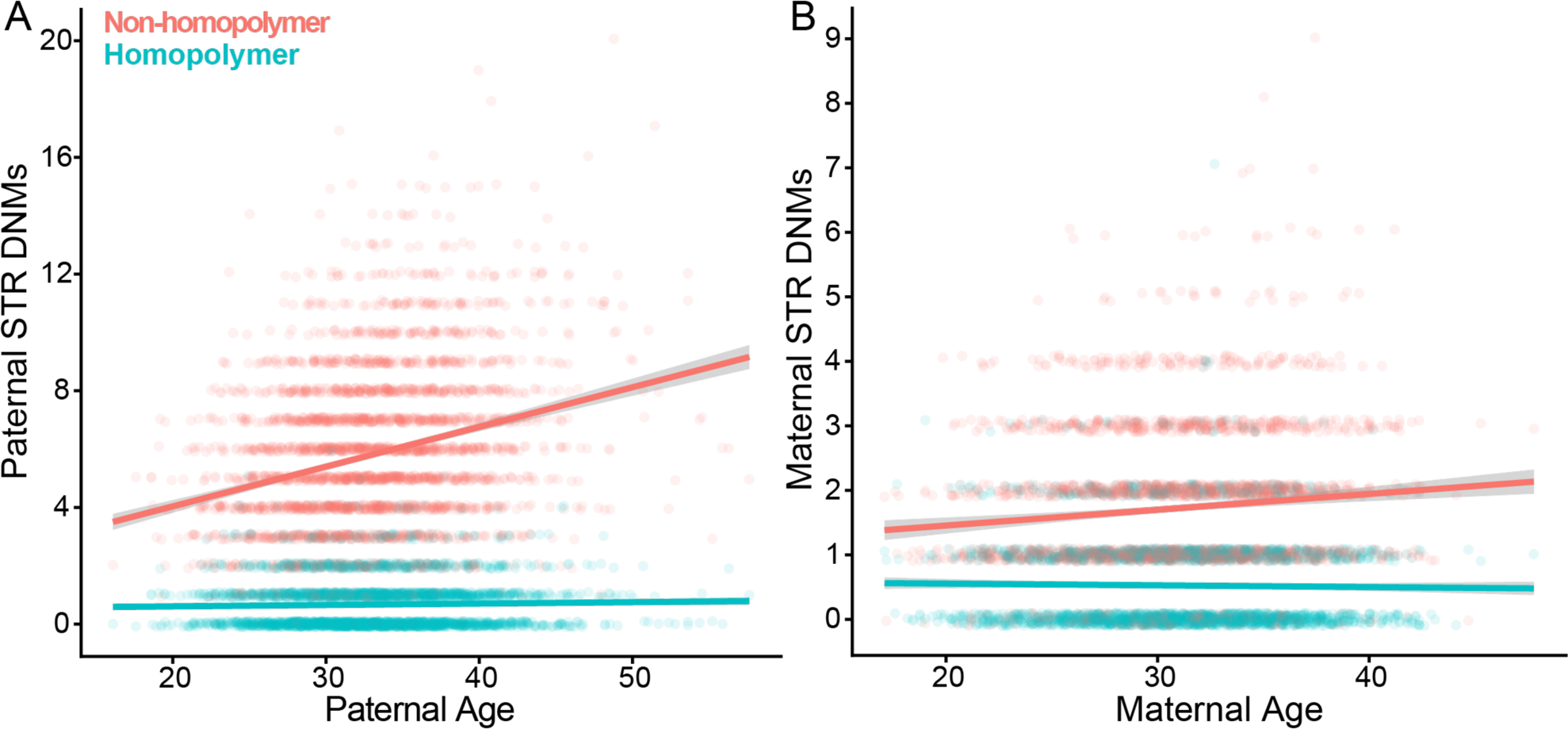
Homopolymers do not harbor parental age effects. We plotted the number of STR DNMs for each trio in the SSC as a function of their parental lineage (paternal vs. maternal) and length of repeat unit (homopolymer vs. nonhomopolymer). Poisson GLMs with identity link functions were fit using ggplot2. Homopolymers have no significant association with either paternal or maternal age. Y-axis jitter was added for readability.

Prior to Kristmundsdottir et al. 2023, studies found no significant maternal age effect on the STR DNM rate, but most have used several orders of magnitude fewer loci and DNMs (Sun *et al*. 2012; Forster *et al*. 2015). To determine if these prior studies were simply underpowered to discover the maternal age effect, we repeated our regression-based inference of the DNM rate after restricting to the subset of loci analyzed in a prior study by Sun, et al. that found no maternal age effect (Sun *et al*. 2012). Although this study included more trios than ours, they used a far smaller number of loci – a subset of 2477 STR sites from a larger set of ∼5000 (AC)_n_/(TG)_n_ dinucleotide repeats used as markers to infer recombination maps (Kong *et al*. 2002). Using the superset of Kong et al. sites that mapped to loci in our panel (n = 2040), we found significant association of the rate of paternally derived DNMs but no such association of maternally derived DNMs with maternal age (*P* = 2.81*10^−7^, 0.165 for paternal, maternal age effects; linear regression) (Figure S4) (Methods). These results indicated that the low number of loci in prior studies have resulted in insignificant statistical power to discover a maternal age effect.

**Figure S4:**
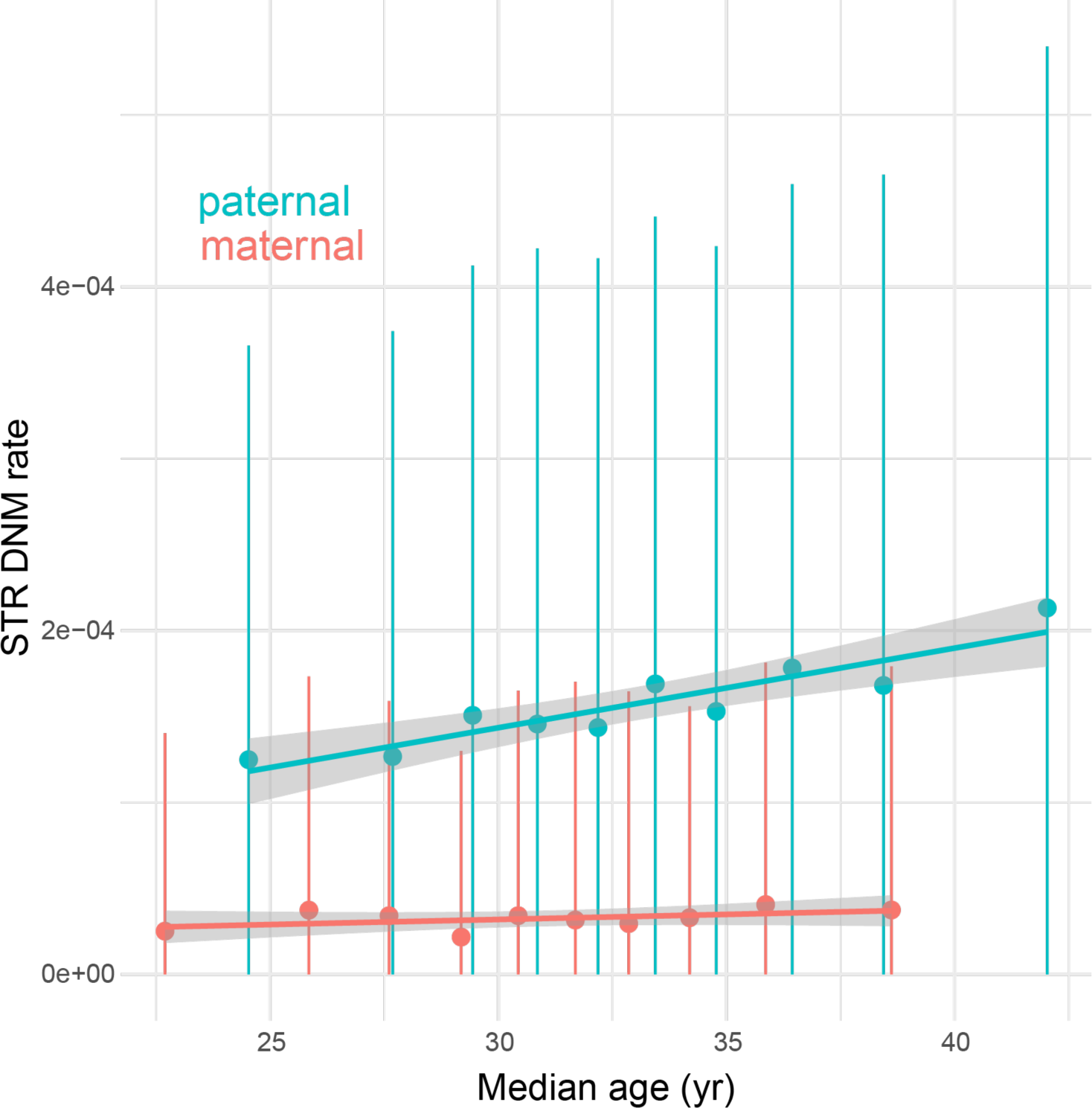
No detectable maternal age effect on loci identified in Kong et al., 2002. Sun et al., 2012 reported no significant maternal age effect on STR DNM. To replicate their analyses in the SSC, we subset our variants to those that overlapped loci from Kong et al., 2002 (N = 2040). We then regressed the maternal mutation rate against maternal age and found no significant association (*P* = 0.165, GLM). The figure above shows the distribution of maternal mutation rates as a function of maternal age deciles; the GLM line was fit with ggplot2.

Recent studies of de novo SNVs have found that the fraction of phased mutations arising from the paternal lineage, known as *ɑ*, does not depend on parental age; instead, the father contributes about 3/4 of mutations regardless of the total mutation load or the ages of the parents (Gao *et al*. 2019). In contrast to this, we find a significant positive association between ɑ and paternal age, indicating that the paternal STR mutation rate increases disproportionately to the maternal mutation rate (0.0023 increase in paternal fraction per year, *P* = 4.99*10^−3^, quasibinomial GLM with identity link function) (Figure S5). This observation likely supports replication as the dominant mechanism of STR DNMs, though the weak slope and nonzero maternal age effect indicate that damage independent of replication still contributes.

**Figure S5:**
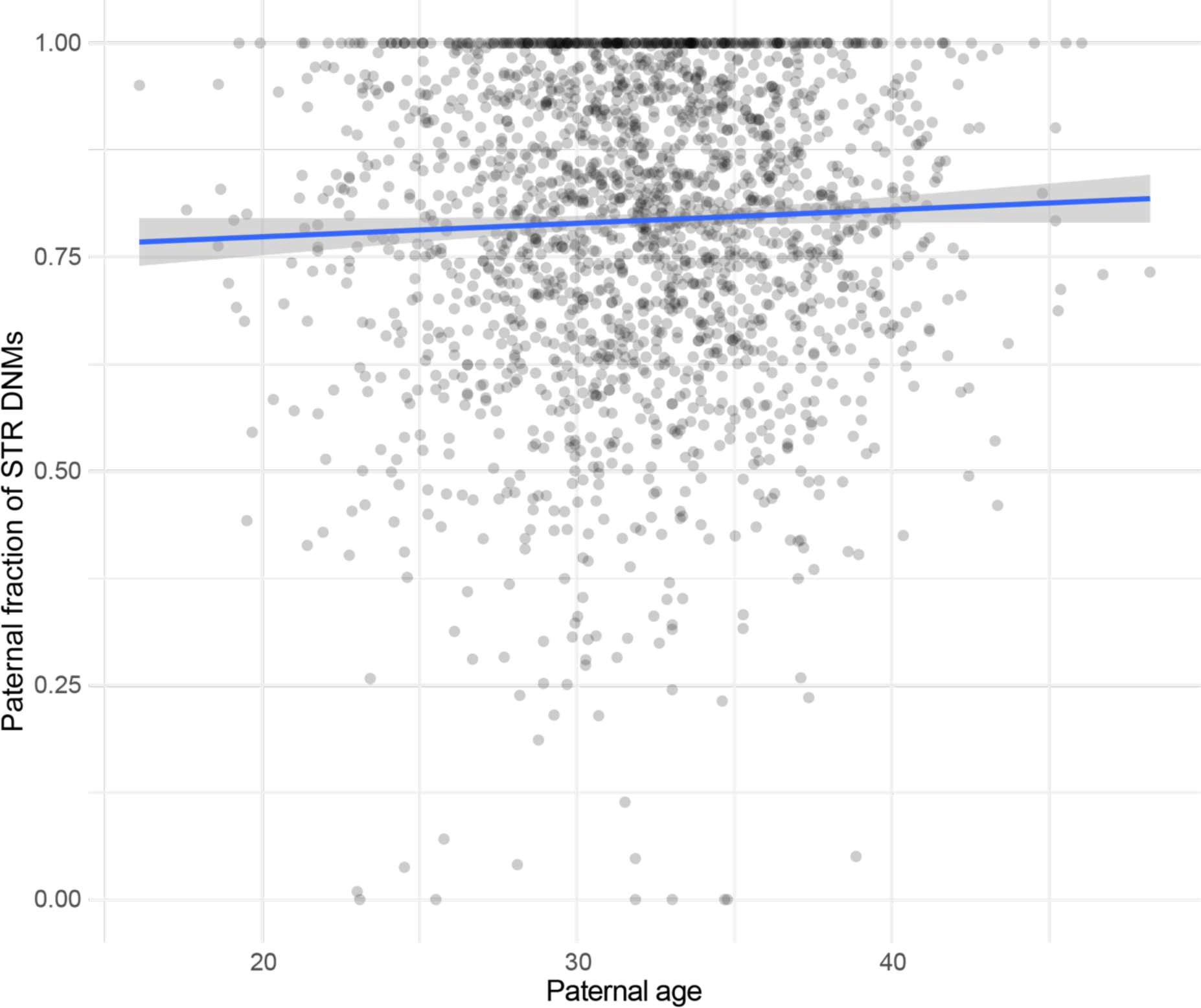
Positive association of paternal age with paternal fraction of mutations. We plotted alpha, the fraction of paternally phased mutations of all phased mutations per trio in the SSC. Quasibinomial regression line with identity link fit with ggplot2. Y-axis jitter was added for readability.

Part of the maternal age effect on single nucleotide substitutions has been attributed to a higher rate of post-zygotic mutagenesis in older mothers (Gao *et al*. 2019). To examine whether the same might be true for STRs, we tested for covariance between maternal age and the mutation rate of STRs occurring on paternally inherited chromosomes, controlling for variance in paternal age (Gao *et al*. 2019). After controlling for paternal age, any correlation of mutation rate on paternally inherited chromosomes with maternal age could indicate that older mothers harbor a higher postzygotic mutation rate. To do this, we randomly sampled nonoverlapping pairs of children born to fathers of the same age (+/− 6 months). For each pair of children, we calculated the differences in maternal ages and the additional number of STR DNMs phased to the father identified in the child born to the older mother. We detected no significant correlation between maternal age and paternally phased STR DNMs, conditional on equal paternal age (one-sided Spearman’s correlation test, *P* = 0.47; GLM *P* = 0.831) (Figure S6). This analysis provided no evidence that advanced maternal age causes additional post-zygotic mutations to accumulate on paternally inherited chromosomes. Simulations, however, suggest that our power to detect weak postzygotic maternal age effects is limited (Figure S7) (Methods).

**Figure S6:**
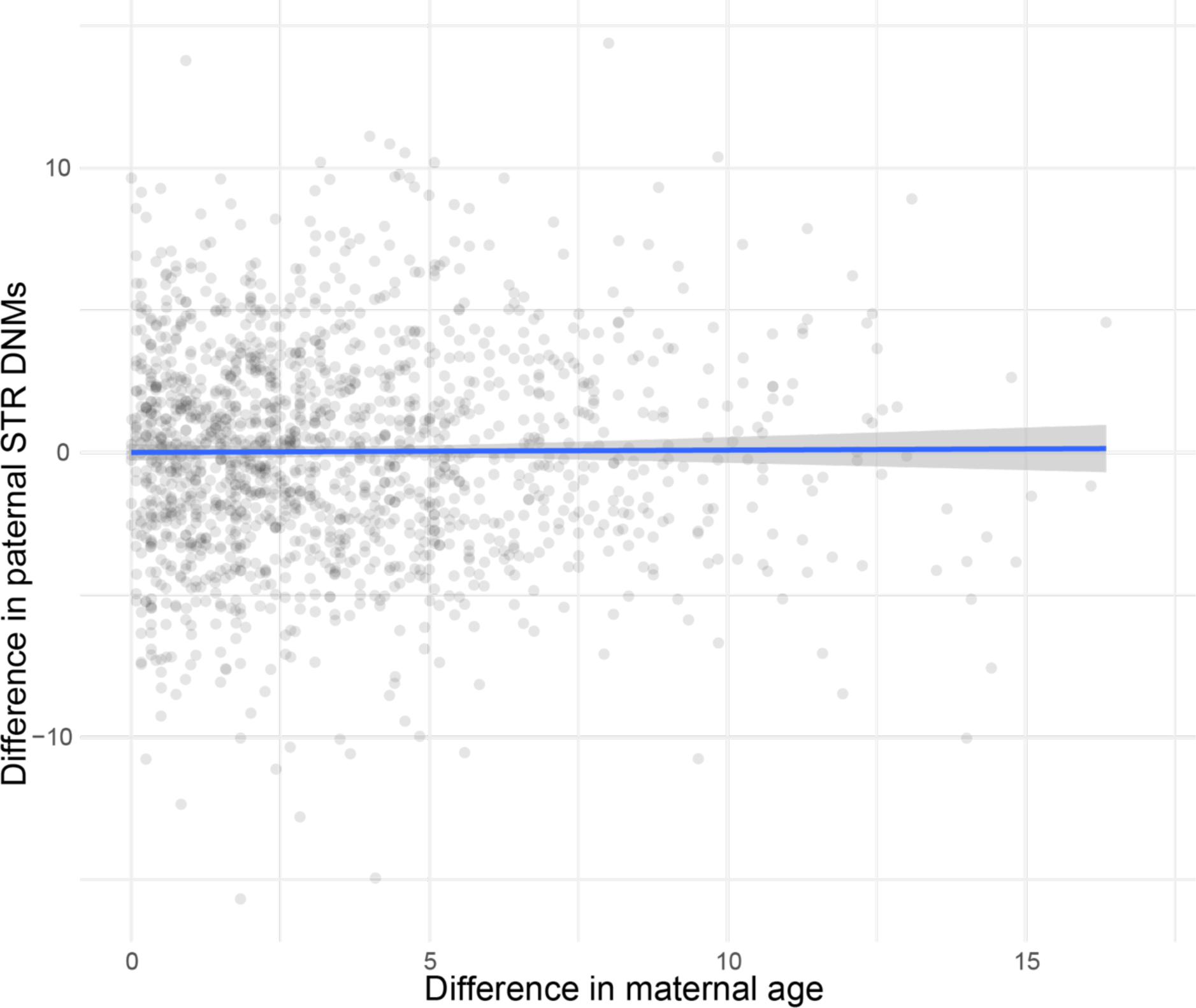
No evidence for postzygotic effect of maternal age on STR DNM rate. Nonoverlapping pairs of children were grouped together by identical (+/− 6 months) paternal ages. Each pair is represented a single time in the scatterplot above, where the difference in the number of paternally derived STR DNMs between the kids is plotted as a function of the difference in maternal age (older – younger). GLM regression line fit with ggplot2, though the slope is not significantly different from zero. X-axis jitter was added for readability.

**Figure S7:**
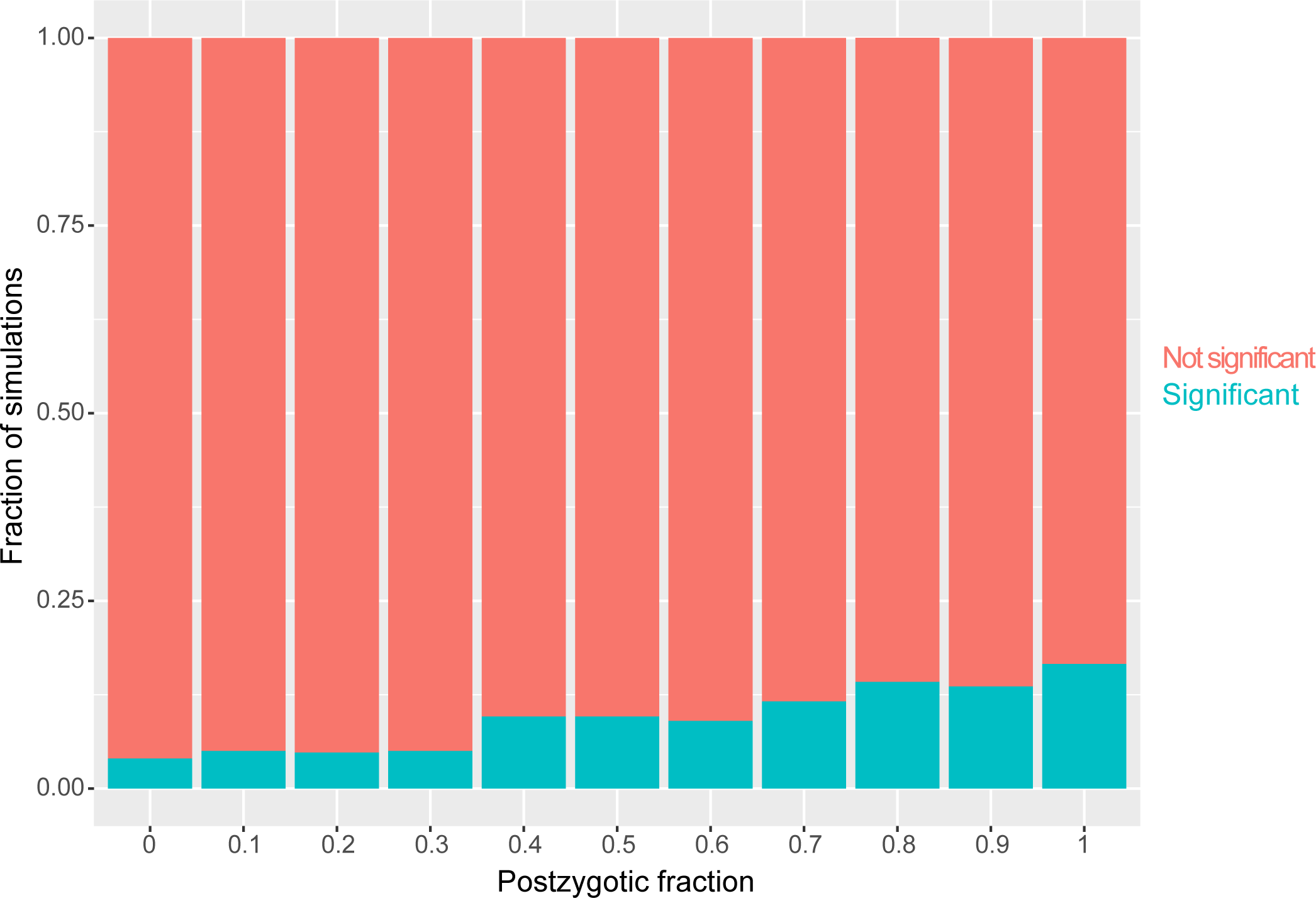
Limited power to detect maternal age effect on postzygotic STR DNMs. The postzygotic fraction represents the fraction of STR DNMs associated with the maternal age effect that occur after zygote formation and are therefore randomly distributed between the paternally and maternally inherited chromosomes. We simulated paternally derived STR DNMs with a variety of postzygotic fractions for each child in the SSC. We then determined the correlation between these (simulated) paternally derived STR DNMs and the difference in maternal age between pairs of children born to fathers of the same ages, as in Figure S6. Plotted are the fraction of *P* values from a one-sided Spearman’s rank correlation test that are significant to *P* < 0.05 from 500 simulations at each postzygotic fraction.

### Nucleotide content modifies parental age and sex effects on STR DNM rate

Thorough characterization of the maternal age effect on SNV DNM rate found that the effect was enriched for specific genomic hotspots and C>G mutations (Jónsson *et al*. 2017; Gao *et al*. 2019). These attributes help suggest that maternal age effect could be partially explained by repair of double-strand DNA breaks. Similarly, we tested whether the STR maternal age effect was biased towards genomic hotspots or characteristic loci.

Certain mutational pathways display biases towards expansions or deletions (Alexandrov *et al*. 2020). Deletion mutations (loss of repeat unit(s)) are less common than expansion mutations (gain of repeat unit(s)) only in the maternal lineage, possibly indicating sex-specific differential mutational pathway activity (Mitra *et al*. 2021). However, we observed no significant difference in parental age effect between deletion and expansion mutations (*P* = 0.54287, 0.212928 for y-intercepts, 0.89, 0.67 for slopes, for paternal and maternal lineages, respectively) (Figure S8).

**Figure S8:**
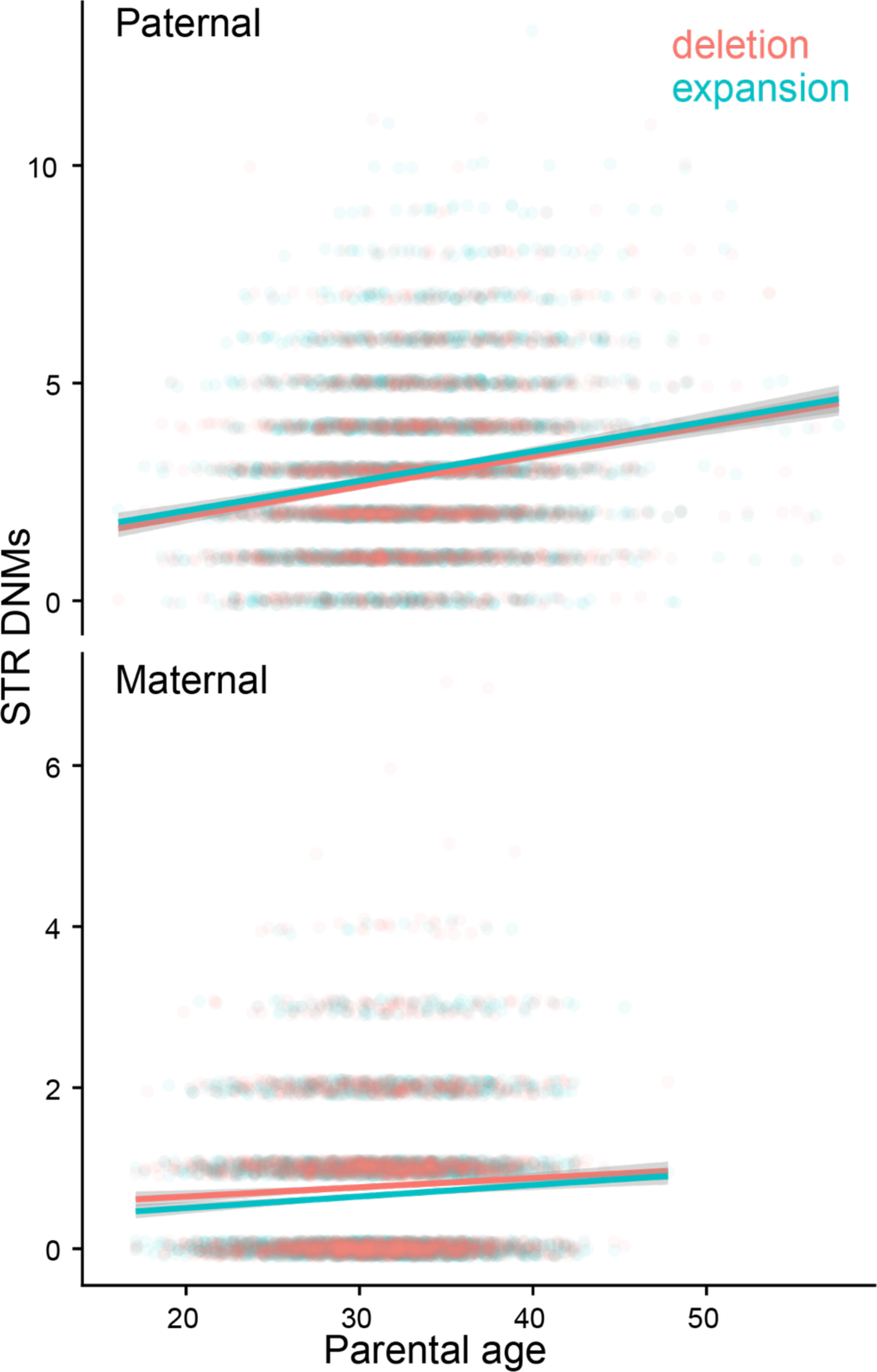
No parental age effects specific to mutation direction. Each trio in the SSC is represented twice on each scatterplot, where the points represent the number of paternal or maternally derived deletion or expansion STR DNMs. Poisson GLMs with an identity link function fit with ggplot2. The slope coefficient of paternal and maternal age is not significantly different between expansions and deletions. Vertical jitter added for readability.

A set of genomic regions in humans have been classified as maternal age hotspots because they have exceptionally elevated mutation rates in the children of older mothers with a particular enrichment of C>G mutations (Jónsson *et al*. 2017). These regions appear to be hotspots for double-stranded breaks in older oocytes, leading to damage-associated mutagenesis. However, we detected no significant association between maternal age and the number of maternally derived STR DNMs (*P* = 0.99, 0.99, Poisson GLM with identity link and Pearson’s correlation test, respectively). These analyses, however, could be underpowered due to the relatively small number of STRs that both passed our filters and occurred in maternal hotspots (N = 432).

Kristmundsdottir et al. previously found that maternal and paternal age affect the DNM rate of STRs differently as a function of their nucleotide composition (Kristmundsdottir *et al*. 2023). We separated STRs into two categories: STRs composed entirely of A/T nucleotides and those including any G/C nucleotides; these categories were chosen to maximize statistical power (Figure 2a). We then modeled the effect of parental age on STR DNM rates, testing for covariance and interaction with STR nucleotide content. To account for differences in the number of loci between nucleotide content bins, we included an offset variable of the number of genotyped nonhomopolymer sites, which forces a beta coefficient of 1 (though using an offset is more straightforward to apply with a log link function, which we used in this figure’s analyses) (Figure 2a). Paternal mutation rate at GC-containing STRs was significantly higher than that of AT-only STRs, though the paternal age effect was not significantly different; in fact, a model without an interaction variable was a better fit while accounting for differences in degrees of freedom (Figure 2b) (*P* < 2 * 10^−16^ for GC content effect on intercept; delta AIC = −2). However, maternal age affected the rate of STR DNMs at A/T-only STRs significantly more strongly than that at G/C-containing STRs (Figure 2c) (*P* = 0.022 for interaction between maternal age and GC content). A model without the interaction variable fit significantly worse, even while accounting for fewer degrees of freedom (delta AIC = −3). Maternal age does not significantly covary with the rate of phasing of either AT-only or GC-containing STR DNMs (*P* = 0.163, 0.893, respectively; binomial regression).

**Figure 2.**
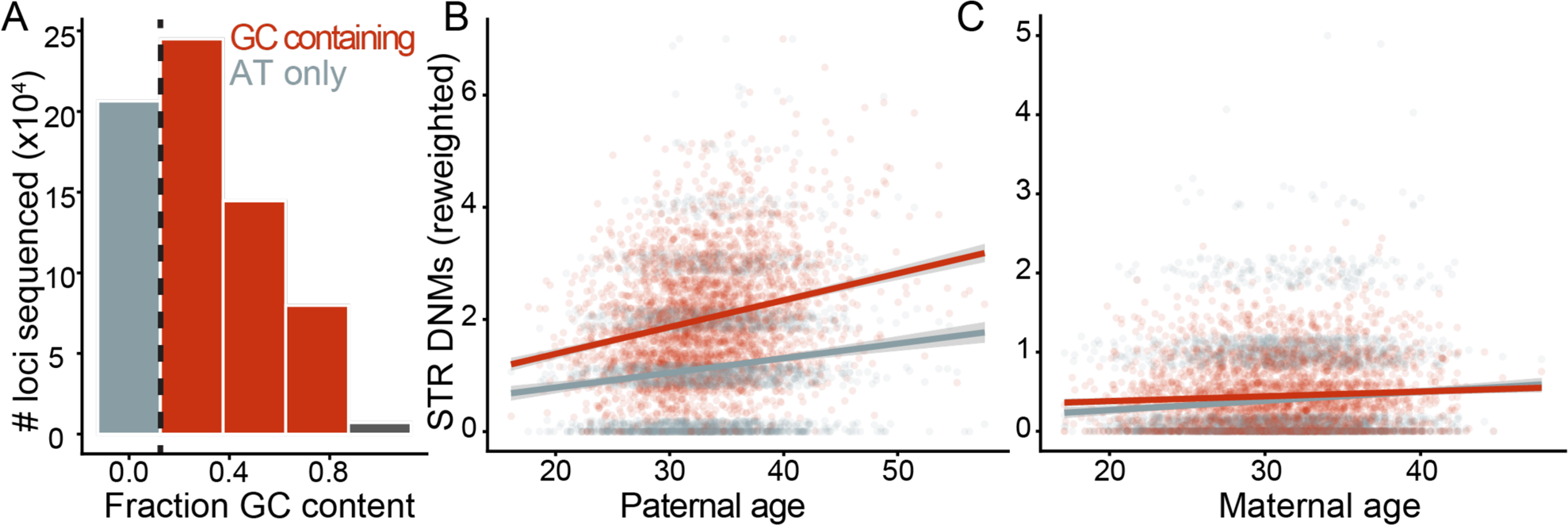
Nucleotide content-specific effects of maternal age. A: We split loci into two bins as a function of their nucleotide content: AT-only and GC-containing. B & C: Association of paternal and maternal age with STR DNM rate as a function of nucleotide content. In each scatterplot, each trio in the SSC is represented twice. The number of STR DNMs phased to the respective parental lineage is plotted as a function of the respective parental age for AT-only and GC-containing STRs. For readability and to account for differences in panel size, the number of GC-containing STR DNMs is rescaled by the ratio of AT-only to GC-containing loci in the panel. We account for panel size differences in the models described in the main text with an offset covariate, though lines plotted here are Poisson GLMs with identity link functions fit to rescaled mutation counts with ggplot2. Coefficients are significant beyond *P* = 0.05.

Certain motifs can form non-B DNA conformations that affect local mutation rate; we therefore tested the maternal age effect on specific unique motifs and their predicted conformations (Guiblet *et al*. 2018; McGinty and Sunyaev 2023). We classified STR loci by their propensity to form H-DNA and Z-DNA (poly purine or poly pyrimidine and purine-pyrimidine with G on one strand, respectively). Z-DNA motifs did not demonstrate a significantly different maternal age effect (*P* = 0.81, Poisson GLM with offset), although STRs likely to form Z-DNA had a significantly higher mutation rate than non-Z-DNA loci (y intercept, *P* = 6.64*10^−15^). A model testing for an interaction between H-DNA formation and maternal age demonstrated that the mutation rate at H-DNA motifs was significantly less affected by maternal age than the rate at non-H-DNA motifs, but the model’s fit was inferior to a simpler model ignoring DNA shape (*P* = 0.014, Poisson GLM with offset; delta AIC = −1918). After collapsing STR motifs by periodicity and reverse complementation (i.e., grouping together loci with AC, CA, GT, and TG motifs), we estimated the maternal age effect on STRs of each common motif separately (> 20 DNMs called that passed filters). The two motifs that harbored significant maternal age effects were AC and AAAT motifs (*P* < 0.05, Bonferroni correction); neither motif is self-complementary and therefore not predicted to commonly form hairpins (Guiblet *et al*. 2018).

In summary, we found that nucleotide content modifies the maternal age effect on STR DNMs. Few other *cis*-acting attributes similarly modify parental age effects; the consistency of the maternal age effect across different variables supports its validity.

### No significant effects of parental age or sex on distribution of fitness effects of STR DNMs

Prior research on this cohort has shown that probands diagnosed with ASD harbor more DNMs at STRs than their unaffected siblings and that these mutations are predicted to be more deleterious (Mitra *et al*. 2021). This evidence supports claims that ASD may be somewhat related to DNM burden at STRs. Following our finding that maternal age more strongly affected certain types of STRs, we sought to test whether parental age and sex affected the deleteriousness of DNMs. Selection coefficients were inferred for ∼90,000 di-, tri-, and tetranucleotide STRs using SISTR as described in (Mitra *et al*. 2021). Briefly, SISTR infers parameter *s* by fitting allele frequencies observed in the unaffected parents at each site to a model that incorporates mutation, demographic history, and selection. This parameter refers to the increase in the selection coefficient (i.e. decrease in reproductive fitness) associated with each additional repeat unit away in length from the major allele, assumed here to be the most fit. As in Mitra et al., we excluded sites for which the posterior distribution of *s* was greater than 0.3, indicating poor model fit; this left ∼63,000 sites. Notably, our additional filters accounting for possible allelic dropout removed 38.1% (20,419 of 53,602) STR DNMs for which a selection coefficient could be inferred.

To initially test for parental age effects on the distribution of fitness effects (DFE) of STR DNMs, we aggregated the allele-specific selection coefficients of paternally and maternally derived mutations across all children born to parents below and above the median age for each sex, respectively. We found no significant differences in the DFE (two-sided Kolmogorov-Smirnov test, *P* = 0.7487, 0.292, respectively). We also modeled the effects of parental age and sex on DFE with Poisson models regressing the number of paternal and maternal STR DNMs at constrained or neutral sites as a function of parental age. Neutral loci were those with an *s* = 0; constrained loci had an *s* in the upper 90^th^ percentile of all genotyped loci with *s* values. In these Poisson models, we tested for linear interaction between constraint and parental age while accounting for differences in the number of loci with an offset variable. The interaction terms were not significant for either parental sex, indicating no significant effect of parental age or sex on DFE (*P* = 0.459, 0.21591). Furthermore, in contrast to Mitra et al., we did not observe any significant differences in the DFE between STR DNMs found in probands and unaffected siblings in the database (two-sided Kolmogorov-Smirnov test on selection coefficients summed across all children by diagnosis, *P* = 0.515). This difference may simply reflect the lower statistical power associated with our strict filtering methods, as described above.

## Discussion

Our analyses of the parental age effects on STR mutagenesis support the hypothesis that STR mutagenesis is more replication dependent than SNV mutagenesis, but do not support the dogma that replication slippage during S phase is the sole cause of STR mutations. The fraction of phased mutations deriving from the paternal lineage increases with paternal age, supporting a hypothesis that the ratio of paternal to maternal mutation rates is lower pre-puberty than post-puberty. This is ostensibly a result of the continued replication of the paternal germline after puberty. Nevertheless, the significant maternal age effect cannot be a result of polymerase slippage during S-phase replication, as the maternal germline does not replicate after birth (Drost and Lee 1995). This observation is reminiscent of findings of STR expansions in postmitotic cells and may provide broader evidence that damage-associated STR mutagenesis is observed across many cell types (Gonitel *et al*. 2008).

The stronger maternal age association with deletion DNMs at AT-only repeats may provide a clue as to the source of these mutations, though our current understanding of STR mutational signatures is not advanced enough to decode it. A number of indel mutational signatures in COSMIC, a database of signatures deconvoluted from mutation spectra identified in tumors, have proposed etiologies such as TOP1 transcription-associated mutagenesis (Alexandrov *et al*. 2020; Reijns *et al*. 2022). However, no specific indel signature matches well with our observed maternal age effect, enriched at AT repeats but without bias towards expansions and deletions. Future work may further integrate this mutational data with somatic mutational data, particularly from post-mitotic cells, to help elucidate possible pathways.

## Methods

### DNM filtering

We analyzed published STR genotypes and DNMs in the SSC as previously described in Mitra, et al. (2021). However, we implemented several additional filtering strategies to account for possible cryptic sources of bias in the dataset. In particular, we noticed that DNMs at loci where one parent is homozygous and the other is heterozygous are more likely to originate on the chromosome inherited from the homozygous parent rather than the heterozygous parent (Figure S1). We hypothesized that this observation might be the result of allelic dropout events: if a parent is heterozygous at a locus but incorrectly genotyped such that they appear homozygous, their child could inherit an allele that is falsely identified as a de novo event. Similar strategies to account for possible allelic dropout in this dataset have been previously described; we developed a new, strict, read-based approach to accounting for this possible artifact (Mitra *et al*. 2021).

Using a read-based approach, we filtered a starting set of 175,290 *de novo* STR calls identified by MonSTR. We used pysam to assess the number of Illumina reads with the *de novo* allele in a child and its parents. Importantly, we did not perform a local realignment step like Mitra et al., but instead opted to use the original alignments to GRCh38. First, we selected any primary reads aligned to the starting coordinate of the STR with mapping quality > 60. We retained only reads that spanned not only the length of the longest observed STR allele in the family, but also included 10 base pairs of flanking sequence on either end of the STR, excluding soft-clipped bases. For every read that completely spanned the length of the longest STR sequence and its flanking bases, we queried the sequence at those coordinates. To determine the length of the STR allele in a given read, we simply traversed the sequence until we found the first instance of the STR motif, and then counted the number of continuous STR motifs that followed. Once we reached the end of the STR, we checked the 10 bases preceding the start and following the end to ensure that they matched the reference sequence that flanks the STR. Reads with interruptions in the STR sequence or more than one mismatch between the observed and reference flanking were discarded. If there were no reads with the *de novo* allele in the child, we called the variants a false positive. If there was one or more reads with the *de novo* allele in either parent, we called the variant inherited. All *de novo* variants with no read support in either parent and at least one supporting allele in the child were labeled true events, for a final *de novo* STR callset of 56,925 variants.

We generated long read sequencing data using Pacific Biosciences high-fidelity (HiFi) and Oxford Nanopore Technologies (ONT) for two families from our dataset in order to further validate *de novo* variants and assess our validation strategy (a total of 157 STR variants across 4 children) (Table S1). For the parents and child, we aligned HiFi and ONT data to GRCh38, and then manually examined the length of the STR allele in each aligned read with mapping quality > 60. Using the same criteria (at least one read with the *de novo* allele in the child, and no reads with the *de novo* allele in the parents), we determined whether each variant was a true de novo event. Upon visual inspection, we found that only 52.5% (31/59) of non-homopolymer DNMs have no evidence of being inherited (24.8% of all DNMs [39/157]). Furthermore, we found that only 8% of putative STR DNMs at homopolymer loci showed no evidence of being inherited from parents or falsely genotyped in the child. This observation further motivated our decision to exclude all homopolymer DNMs from analysis. Based on the long read data, we predict our filter has an accuracy of 92.3%, sensitivity of 94.8%, and specificity of 91.5%. Furthermore, the bias towards DNM phasing to homozygous parents we observed with the full dataset was attenuated when applying these filters (Figure S1).

The sensitivity of this filtration strategy, however, may be lower at some loci and alleles. Each read mapping to an STR locus supports a certain allele length, here measured in number of repeat units; GangSTR uses the amount of read support for allele lengths to call a genotype. However, at some loci, reads support a variety of possible allele lengths. For example, at a site where parents have genotypes [10, 10] and [10, 10] but frequently we observe reads supporting an allele of length [11], we would erroneously filter possible *de novo* [11] alleles in a child. We were therefore motivated to further assess the variance in sensitivity of our filter, i.e., how frequently would we erroneously filter out a putative STR DNM. Given a family with parental genotypes [10, 10] and [10, 10] and a proband genotype of [10, 11] (putative *de novo* allele [11]), we searched for any other set of parents with the exact same genotypes (sex non-specific) and no children with the same putative *de novo* allele; these are “positive” couples, from whom a putative [11] allele was unlikely to have dropped out from genotyping. In these positive couples, we counted the number of reads that supported the target family’s putative *de novo* allele to assess the frequency with which we would erroneously flag a couple for possible allelic dropout. We found that sensitivity differed greatly between loci and sometimes between alleles at the same locus (Figure S2). We could not reliably predict which loci and genotypes had high false negative rates; calculating the empirical false negative rate was only possible for loci and genotypes common among the SSC parents. Implementing an updated filtration strategy allowing >0 reads supporting the putative *de novo* allele at certain loci could bias us against rare genotypes, especially those uncommon in families of predominantly European ancestry, which comprise the bulk of the SSC (Wilfert *et al*. 2021). Thus, we chose to maintain our strict filtering technique and accounted for possible variance in sensitivity by calculating and regressing with an accurate denominator, as described in the following section.

### Denominator calculation

At sites with high sequencing noise or parental diversity in alleles, only a subset of possible DNMs can be confidently called. In the following descriptions, alleles are annotated by their number of repeat units. For example, at a site where the parents have genotypes [10, 10] and [11, 11], a 10 -> 11 mutation or an 11 -> 10 DNM resulting in either a [10, 10] or a [11, 11] genotype in the child would be indistinguishable from (and perhaps more parsimoniously explained by) allelic dropout in the child. Furthermore, if ≥ 1 reads from the [11,11] parent indicated a 12 allele, observing the 12 allele in the child could again indicate dropout over mutation. However, any DNM could be discoverable at a site where both parents are [10, 10] with no reads mapping any other alleles. These concerns affect the denominator for each trio, i.e. how many sites are we able to discover DNMs. To account for variance in denominator as a possible artifactual source of the association of STR DNM rate with maternal age, we applied an extra set of filters requiring 100% discoverability for all DNMs observed in a trio. For a given mutation size of *i* repeat units, a site is fully discoverable if and only if for every parental allele *a* in all parental alleles *A*, no reads in the parents map to any allele of size *a +/− i*. Note that this filter excludes possible DNMs of size *i* that occur at sites where not all DNMs of size *i* are discoverable. The resulting mutation rate is calculated as

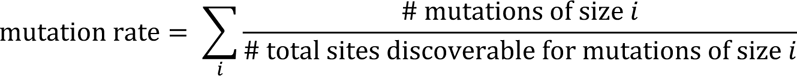

We calculated the mutation rate across *i* in [1, 2, 3], which accounted for 98% of the DNMs which passed filters described in the previous section.

To implement this filter, we used the same underlying framework of our filtering script to identify STR sites where we could observe a de novo mutation of a given number of motifs. For each pair of parents in the dataset, we examined every STR site that was genotyped in all four parental haplotypes in Illumina data aligned to GRCh38. Borrowing code from our STR filtering script, we selected passing reads and estimated the length of the STR in each read. We classified every observed allele as an “uncallable mutation” – if an allele was observed in even 1 parental read, it would never pass our *de novo* filtering script. Unfortunately, this filter left us with only 833 maternally derived mutations, which gave us insufficient power to observe a significant maternal age effect on their rate per child (*P* = 0.12, GLM). These validated mutations are more challenging to phase with our methods, given the high proportion from sites with no diversity in parental alleles (31% vs. 15% of DNMs that passed filters in prior section). However, a regression of the mutation rate of all mutations regardless of phase and against both maternal and paternal age found that both parental ages were significantly positively associated with rate (*P* = 1 * 10^−11^, 0.012 for paternal, maternal age). This model was significantly better than one that did not include maternal age as a covariate (delta AIC = −4). Furthermore, Poisson GLMs regressing the number of fully discoverable mutations (numerator in above equation) against parental age including an offset of the sum of the number of loci discoverable for mutations of sizes 1, 2, and 3 (denominator in above equation) found significant paternal and maternal age effects, respectively (*P* < 2.2 * 10^−16^, *P* = 4.41 * 10^−9^ for paternal and maternal age effects, respectively, log link). This indicated to us that, although the maternal age effect appeared robust to differences in denominator, this approach could result in a lack of statistical power for any further analyses. Therefore, in the main figures, we continued to use the filtering approach described in the previous section that did not account for discoverability of sites.

### Determining power to detect postzygotic effect of maternal age

We find no evidence that the maternal age effect on STR DNMs is due to a heightened amount of postzygotic mutagenesis in the child’s early development, in contrast to what has been detected in SNV DNMs (Gao *et al*. 2019). However, due to the low number of nonhomopolymer STR DNMs that pass our filters and can be phased to a parental lineage, we may be limited in our power to detect small amounts of this postzygotic mutagenesis. To determine our power, we simulated paternally derived mutations for children in the SSC at a variety of postzygotic maternal age effects and tested for correlation between maternal age and paternally derived DNMs while conditioning on paternal age, as in Figure S6. If some fraction *f_pz* of the additional mutations associated with maternal age occur after zygote formation, we assume that these mutations will be randomly distributed between the paternally and maternally inherited chromosomes. Thus, given paternal and maternal ages *age_pat_* and *age_mat_* per child, paternal age y-intercept and slope *B_0,pat_* and *B_1,pat_*, and maternal age slope *B_1,mat_*, we simulated paternally derived STR DNMs as a Poisson distributed random variable given by λ:

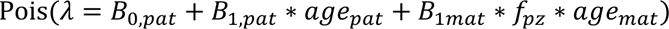

Beta coefficients were taken from the Poisson linear models in Figure 1. Poisson sampling was done in R 4.1.1 using function `rpois()`. For each round of simulation, we drew paternally derived STR DNMs for each child, grouped children into nonoverlapping pairs of identical rounded paternal ages, and calculated the difference in maternal ages and the difference in simulated paternally derived STR DNMs between the child born to younger and the child born to older parents. We then estimated the significance of positive correlation between the maternal age difference and paternal STR DNM difference for all pairs of children with a one-sided Spearman’s rank correlation coefficient and saved the *P* value. For values of *f_pz_* < 0.4, we appear to have no power to detect significant postzygotic effects; our power is still limited even for stronger postzygotic effects. More efficient phasing methods, cleaner genotyping, and a larger set of families would all increase statistical power for future iterations of this analysis.

### Regression framework

To model the effects of parental age on the number of STR DNMs, we typically employed Poisson regressions with identity link function in R (version 4.1.1), following prior work (Jónsson *et al*. 2017; Sasani *et al*. 2019). The identity link specifies a linear relationship between the parental age and the expected number of mutations in a given child at that age. For example, we used the following model to estimate the maternal age effect on number of maternally-derived non-homopolymer STR DNMs.

~~~
glm(maternal_nonhomopolymer ∼ motherAgeAtBirth, … family = poisson(link = “identity”))
~~~

We similarly modeled the effects of nucleotide content and parental age as a Poisson GLM while allowing linear interaction between parental age nucleotide content of STRs (Figure 2). To account for differences in the number of AT-only and GC-containing non-homopolymers in our panel, we included an offset term which directly accounts for differences in denominators for rates, though are more straightforward to implement with a log rather than an identity link function. For example, we used the following model to estimate how nucleotide content modified the maternal age effect:

~~~
glm(n_muts ∼ motherAgeAtBirth + gc_content + offset(log(denom)…, family = poisson(link = “log”))
~~~

In the above model, gc_content is a binary variable classifying a locus as being AT-only or GC-containing; denom is the number of loci in the panel matching the gc_content classification.

### Comparison to Sun et al., 2012

Prior research has detected no significant maternal age effect on the rate of maternally derived STR DNMs (Sun *et al*. 2012). We hypothesized that this study was underpowered to discover the maternal age effect we observed. To test this hypothesis, we subset our loci roughly to the ones that were included in the prior study. Sun et al. examined 2477 loci of a subset of 5136 loci originally identified to estimate a human genetic map (Kong *et al*. 2002; Sun *et al*. 2012). These loci are all (AC)_n_/(TG)_n_ dinucleotide repeats; their discovery process is described in several earlier publications (Weissenbach *et al*. 1992; Dib *et al*. 1996). The list of 2477 loci is not available, so we analyzed the superset of 5136 loci. We found the ∼100-400bp genomic region in GRCh38 associated with the original marker using tables from the UCSC Table Browser. 2040 AC (CA, GT, or TG) repeats in our panel that fell within these windows were included in our analysis. GLMs regressing paternal or maternal mutation rate against paternal or maternal age, respectively, found a significant paternal but no significant maternal age effect (*P* = 2.81*10^−7^, 0.165, respectively). A possible explanation for the lack of significant maternal age effect could be the lower number of transmissions observed at AC dinucleotides in Sun et al.. While their study sequenced 24832 trios at 2477 loci for a total of 6.15 * 10^7^ transmissions at AC dinucleotides, our study examined 3172 at 55830 AC dinucleotide loci alone for 1.77 * 10^8^ transmissions (and 6.83*10^5^ non-homopolymer STR loci for 2.17 * 10^9^ total transmissions).

## Acknowledgements

This work was supported by the National Institute of General Medical Science (T32GM081062 to M.E.G., 1R35GM133428-01 to K.H.), the National Human Genome Research Institute (T32HG008962 to M.E.G.), and the Eunice Kennedy Shriver National Institute of Child Health and Human Development (R01HD106112 to A.R.Q.); the National Institute of Mental Health (R01MH101221 to E.E.E.), all at the National Institutes of Health; the Simons Foundation (SFARI 810018EE to E.E.E.); the Burroughs Wellcome Fund (a Career Award at the Scientific Interface to K.H.); the Pew Charitable Trusts (Biomedical Scholarship to K.H.), the Searle Scholars Program (Career Award to K.H.), and the Alfred P. Sloan Foundation (Research Fellowship to K.H). The authors thank Harriet Dashnow, Ziyue Gao, Priya Moorjani, Phil Green, Sharon Browning, Brian Browning, William Noble, Rick McLaughlin, Melissa Gymrek, and members of the Harris, Eichler, and Quinlan labs for helpful discussions. They also thank Amy Wilfert for technical assistance.

## Code and Data Availability Statement

Code to generate figures and run statistical tests is available on Github at https://github.com/goldmich/str_dnm. VCFs and STR DNMs published in Mitra et al. 2021 are available through SFARI to approved researchers at SFARI Base with base accession code: SFARI_SSC_WGS_2b. Phenotype and sequencing data for the SSC are also available through SFARI Base (accession nos. SFARI_SSC_WGS_p, SFARI_SSC_WGS_1, and SFARI_SSC_WGS_2). HiFi and ONT sequencing data for two quads have been deposited to SFARI base but under embargo until February 2024. liftOver and associated files for the comparison with Sun et al., 2012 were downloaded from the UCSC table browser (https://genome.ucsc.edu/cgi-bin/hgTables).

